# Activation of dopamine receptor 2 (DRD2) prompts transcriptomic and metabolic plasticity in glioblastoma

**DOI:** 10.1101/454389

**Authors:** Seamus P. Caragher, Jack M. Shireman, Mei Huang, Jason Miska, Shivani Baisiwala, Cheol Hong Park, Miranda R. Saathoff, Louisa Warnke, Ting Xiao, Maciej S. Lesniak, C. David James, Herbert Meltzer, Andrew K. Tryba, Atique U Ahmed

## Abstract

Glioblastoma (GBM) is one of the most aggressive and lethal tumor types. Evidence continues to accrue indicating that the complex relationship between GBM and the brain microenvironment contributes to this malignant phenotype. However, the interaction between GBM and neurotransmitters, signaling molecules involved in neuronal communication, remains incompletely understood. Here we examined, in both sexes of humans and mice, how the monoamine dopamine influences GBM cells. We demonstrate that GBM cells express DRD2, with elevated expression in the glioma-initiating cell (GIC) population. Stimulation of DRD2 caused neuron-like depolarization exclusively in GICs. In addition, long-term activation of DRD2 heightened the sphere-forming capacity of GBM cells as well as tumor engraftment efficiency. Mechanistic investigation revealed that DRD2 signaling activates the hypoxia response and functionally alters metabolism. Finally, we found that GBM cells synthesize and secrete dopamine themselves, suggesting a potential autocrine mechanism. These results identify dopamine signaling as a potential therapeutic target in GBM and further highlight neurotransmitters as a key feature of the pro-tumor microenvironment.

**Significance Statement:** This work offers critical insight into the role of the neurotransmitter dopamine in the progression of GBM. We show that dopamine induces specific changes in the state of tumor cells, augmenting their growth and shifting them to a more stem-cell like state. Further, we show that dopamine can alter the metabolic behavior of GBM cells, increasing glycolysis. Finally, we show that GBM cells, including tumor samples from patients, can synthesize and secrete dopamine, suggesting an autocrine signaling process underlying these results. These results describe a novel connection between neurotransmitters and brain cancer, further highlighting the critical influence of the brain milieu on GBM.

## Introduction

Glioblastoma (GBM) is the most common and aggressive primary malignant brain tumor afflicting adults, with standard of care treatment carrying a median survival of only 15 months(Alifieris and Trafalis 2015). Recent clinical trials have shown increased survival to approximately 21 months(Stupp et al. 2017). A key factor limiting the efficacy of treatment is the remarkable plasticity exhibited by GBM cells, which allows them to effectively adapt to changes in the microenvironment induced by conventional radiation and chemotherapy(Safa et al. 2015; Olmez et al. 2015). Our group and others have shown that cellular plasticity processes enable GBM cells to adopt many phenotypes, including the glioma initiating cell (GIC) state, characterized by expression profiles similar to those of normal neural stem cells and heightened resistance to therapy(Safa et al. 2015; Olmez et al. 2015; Dahan et al. 2014; Lee et al. 2016). Given the dynamic nature of this process, a key question is what mechanisms regulate and control cellular plasticity. Several factors have been shown to activate GBM plasticity, including hypoxia, acidity, and metabolic changes(Hardee et al. 2012; Heddleston et al. 2009; Hjelmeland et al. 2011; Soeda et al. 2009; Tamura et al. 2013). All of these factors are influenced by the dynamic microenvironmental milieu occupied by GBM cells. Therefore, further analysis of the CNS-specific environmental factors affecting tumors are critical for devising novel therapies for patients with GBM.

The GBM microenvironment is unique for a multitude of reasons, including the range of central nervous system (CNS)-specific factors, such as neurotransmitters and neurotrophins, known to influence cell proliferation and differentiation. Given the evidence that plasticity can be induced in GBM tumor cells by a variety of microenvironmental factors, including oxygen availability and acidity(Hjelmeland et al. 2011; Soeda et al. 2009; Evans et al. 2004), it is logical that CNS-specific cues like neurotransmitters may also influence GBM cellular behavior. Indeed, evidence is accumulating that CNS-specific cues contribute to the onset and progression of GBM(Charles et al. 2011). It has been shown, for example, that factors secreted from neurons such as neuroligin-3 directly augment GBM cell growth(Venkatesh et al. 2015; Venkatesh et al. 2017). Further, Villa and colleagues have demonstrated that GBM cells utilize cholesterol secreted by healthy astrocytes to enhance their growth(Villa et al. 2016). In addition, glutamate, a neurotransmitter, has been shown to augment GBM growth via effects on EGFR signaling(Schunemann et al. 2010). Clearly, CNS-enriched factors like neurotransmitters can influence GBM.

One neurotransmitter family likely to contribute to the pro-tumor brain microenvironment in GBM is the monoamines (dopamine, serotonin, and norepinephrine), critical to behavior, emotion, and cognition(Beaulieu and Gainetdinov 2011; Lammel, Lim, and Malenka 2014). A recent review highlighted the potential influence of monoamines on GBM tumors(Caragher et al. 2017). Critically, multiple studies have shown that monoamine signaling influences the behavior and phenotype of healthy neural stem cells (NSCs), which have similar expression profiles and functional attributes to GICs(Singh et al. 2004; Galli et al. 2004). In light of these reports, we set out to investigate how DRD2 activation influences GBM signaling and phenotype.

## Materials and Methods

### Cell Culture

U251 human glioma cell lines were procured from the American Type Culture Collection (Manassas, VA, USA). These cells were cultured in Dulbecco’s Modified Eagle’s Medium (DMEM; HyClone, Thermo Fisher Scientific, San Jose, CA, USA) supplemented with 10% fetal bovine serum (FBS; Atlanta Biologicals, Lawrenceville, GA, USA) and 2% penicillin-streptomycin antibiotic mixture (Cellgro, Herdon, VA, USA; Mediatech, Herdon, VA, USA). Patient-derived xenograft (PDX) glioma specimens (GBM43, GBM12, GBM6, GBM5, and GBM39) were obtained from Dr. C David James at Northwestern University and maintained according to published protocols(Hodgson et al. 2009). Notably, these cells are known to represent different subtypes of GBM. GBM43 and GBM12 are proneural, GBM6 and GBM39 are classical, and GBM5 is mesenchymal, according to the subtypes established by Verhaak(Verhaak et al. 2010).

### Determination of Dopamine Levels

Cell culture supernatant was collected, filtered, and flash frozen. The details of the mass spectrometric/UPLC procedure are described elsewhere(Huang et al. 2014). The LC system (Waters Acquity UPLC, Waters Co., Milford, MA, USA) and a Waters Acquity UPLC HSS T3 1.8 μm column were used for separation. The UPLC system was coupled to a triple-quadrupole mass spectrometer (Thermo TSQ Quantum Ultra, Waltham, MA, USA), using ESI in positive mode. All data were processed by Waters MassLynx 4.0 and Thermo Xcaliber software. Data was acquired and analyzed using Thermo LCquan 2.5 software (Thermo Fisher Scientific, Inc., Waltham, MA, USA).

### Animals

Athymic nude mice (nu/nu; Charles River, Skokie, IL, USA) were housed according to all Institutional Animal Care and Use Committee (IACUC) guidelines and in compliance with all applicable federal and state statutes governing the use of animals for biomedical research. Briefly, animals were housed in shoebox cages with no more than 5 mice per cage in a temperature and humidity-controlled room. Food and water were available ad libitum. A strict 12-hour light-dark cycle was instituted.

### MicroArray

RNA was extracted from samples using RNeasyI kit according to manufacturer’s instructions (Qiagen). 1000ng of RNA were utilized for MicroArray analysis according to manufacturer’s directions (Ilumina). HumanHT12 (48,000 probes, RefSeq plus EST) was utilized for all microarrays. All microarrays were performed in triplicate.

### Glycolytic measurement

GBM 39, 12 and 5 were lifted and adhered at a concentration of 0.5-1.5×10^5^ cells per well to XFe 96 well culture plates (Agilent; Santa Clara, CA) using CellTak tissue adhesive (Corning; Corning, NY). We then performed a Mito-Stress test as described in the manufacturer’s protocol utilizing a Seahorse Xfe96 well extracellular flux analyzer. Briefly, cells were attached and suspended in XF Base medium supplemented with 2mM L-glutamine (Fisher; Hampton, NH). After baseline ECAR reading, glucose was injected to a final concentration of 10mM to determine glucose stimulated ECAR rate. Next, oligomycin (1μM final concentration) was injected to determine maximal glycolytic capacity. Lastly, a competitive inhibitor of glycolysis (2-DG; 2-deoxyglucose) was injected (50mM) to validate ECAR as a measure of glycolytic flux. Data was analyzed using Agilent’s proprietary Wave software.

### Electrophysiology

Whole-cell current-clamp recordings of GBM cells were obtained under visual guidance with a Zeiss Axioscop FS (Zeiss; Jena, Germany); fluorescence was used to target GICs. DRD2 agonist was delivered by superfusion of targeted cells using a Picospritzer device (Parker Hannifan; Cleveland, OH, USA). Recordings were made with a Multiclamp 700B (Molecular Devices; Sunnyvale, CA, USA) and Digidata data acquisition devices (Molecular devices) and digitized at 20,000 samples/s. The baseline membrane potential was corrected for the liquid junction potential calculated using pClamp 10 Software (Molecular Devices). Patch-clamp electrodes were manufactured from filamented borosilicate glass tubes (Clark G150F-4, Warner Instruments). Current-clamp electrodes were filled with an intracellular solution containing the following (in mM): 140 K-gluconate, 1 CaCl2 * 6H2O, 10 EGTA, 2 MgCl2 * 6H2O, 4 Na 2ATP, and 10 HEPES with a resistance of ~ 3MΩ. Cells were recorded in artificial CSF (ACSF) containing the following (in mM): 118 NaCl, 3 KCl, 1.5 CaCl2, 1 MgCl2 * 6H2O, 25 NaHCO3, 1 NaH2PO4, and 30 D-glucose, equilibrated with medical grade carbogen (95% O2 plus 5% CO2, pH= 7.4). Chemicals for ACSF were obtained from Sigma-Aldrich. Cells were submerged under ACSF, saturated with carbogen in recording chambers (flow rate, 12 ml/min). Experiments were performed at 30°C ±0.7°C using a TC-344B Temperature Regulator with an in-line solution heater (Warner Instruments).

### Statistical Analysis

All statistical analyses were performed using the GraphPad Prism Software v4.0 (GraphPad Software; San Diego, CA, USA). In general, data are presented as mean (SD) for continuous variables, and number (percentage) for categorical variables. Differences between two groups were assessed using Student’s t-test or Wilcoxon rank sum test as appropriate. For tumorsphere formation assay a score test for heterogeneity between two groups was used and reported as p-value and Chi-Squared for each group. Difference among multiple groups were evaluated using analysis of variance (ANOVA) with post hoc Tukey’s test, or Mann-Whitney U test followed by Bonferroni correction as appropriate. Survival curves were graphed via Kaplan-Meier method, and compared by log-rank test. All tests were two-sided and P value <0.05 was considered statistically significant.

## Results

### Therapeutic stress induces epigenetic modifications and subsequent increased expression of DRD2

In order to begin our investigation of monoamines in GBM, we analyzed the epigenetic status of their receptors during therapeutic stress. We performed a genome wide ChIP-Seq analysis of PDX GBM43 cells for histone 3 lysine 27 (H3K27) acetylation (ac), a marker of open chromatin and active gene transcription, and H3K27 tri-methylation (me3), a maker of closed chromatin and transcriptional repression. Cells were treated with temozolomide (TMZ, 50uM), equimolar vehicle control DMSO, or radiation (2Gy). ChIP-Seq was performed 96 hours after treatment initiation. Examination of all dopamine, serotonin, and noradrenergic receptors showed that the DRD2 promotor alone possessed enrichment of an H3K27ac mark without any change in H3K27me3 (MACS2 Peak Score: 45.0, fold enrichment: 4.023, P-value < 0.0001) (Figure 1A & B, Extended Data Figure 1-1). Further, analysis of the Cancer Genome Atlas (TCGA) revealed that, of the five dopamine receptors, DRD2 mRNA is present at the highest levels (Extended Data Figure 1-1).

**Figure 1:**
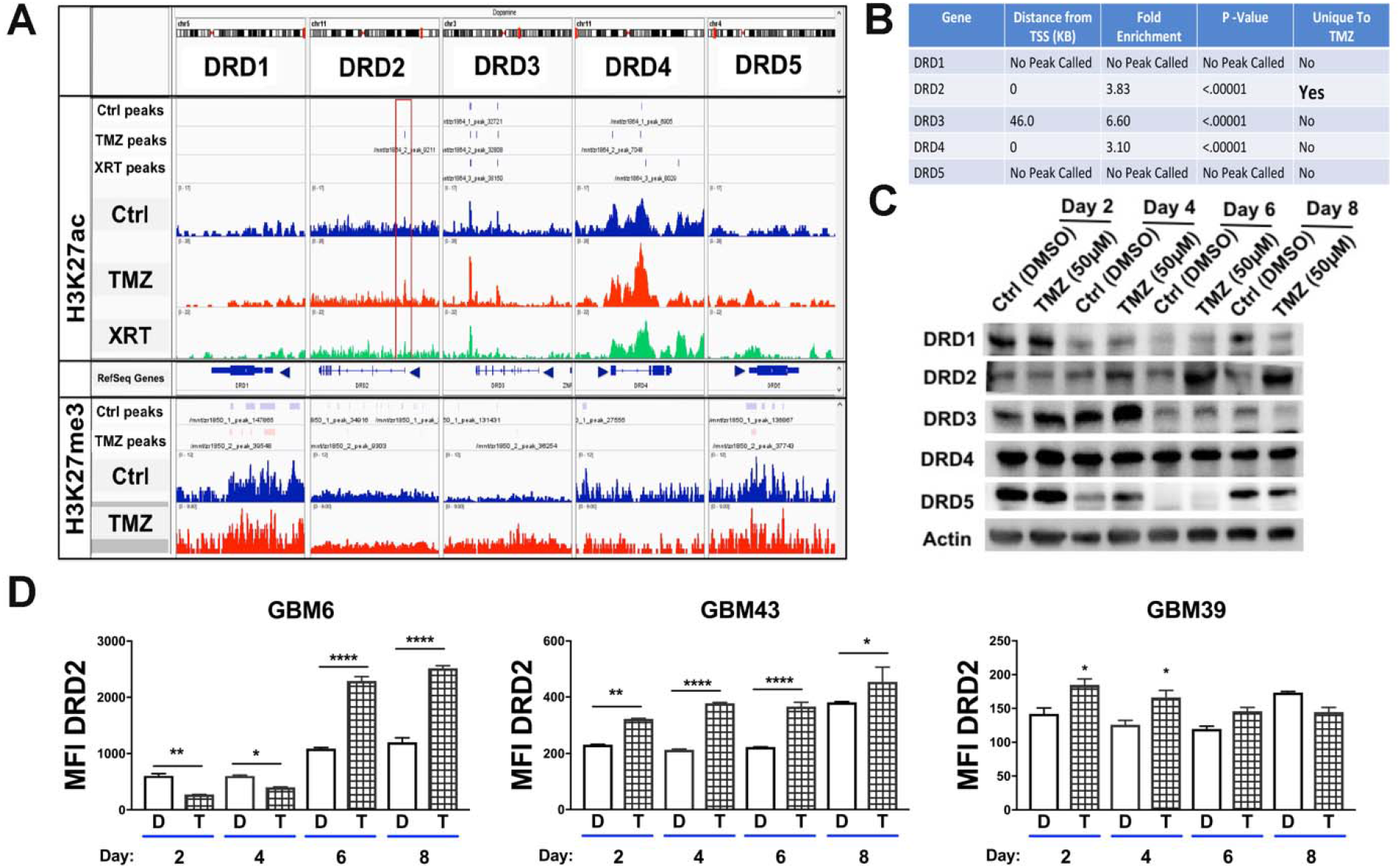
Chemotherapeutic stress induces DRD2 expression in the patient derived xenograft glioma lines. (A/B) H3K27ac and H3K27me3 marks distribution around the transcription start sites of dopamine receptors following therapy. Chromatin immunoprecipitation with massively parallel DNA sequencing (ChIP-Seq) was performed for the open chromatin state histone 3 lysine 27 (H3K27) acetylation in PDX line GBM43 after 4 days of single exposure to temozolomide (TMZ, 50μM), or treated with one fractionated dose of radiotherapy (2 Gy). DMSO was used as a vehicle control. The red box in DRD2 highlights a key residue near the promoter that uniquely exhibited increased H3K27ac following treatment. (MACS2 Peak Score: 45.0, fold enrichment: 4.023, P-value < 0.0001). Table summarizes statistical information for each of the 5 DRDs. Extended data included in Figure 1-1 also shows peak calling of all monoamine receptors. (C) Western blot analysis of dopamine receptors expression in PDX cells treated with TMZ over 8 days. (D) Fluorescence activated cell sorting analysis (FACS) analysis of DRD2 on different subtype of PDX cells treated with vehicle control DMSO or TMZ (50μM). Extended data figure 1-2 includes analysis of all 5 dopamine receptors demonstrating elevated DRD2 expression in human and mouse samples of both sexes. **Bars** represent means from three independent experiments and **error bars** show the standard deviation (SD). Student T-tests were performed for each day separately. *P<.05, **P<.01, ***P<.001.

To determine if this alteration in histone status of the DRD2 promoter leads to increased protein expression, we analyzed protein levels of all DRDs in PDX GBM cells treated with either vehicle control DMSO or TMZ for 2, 4, 6, or 8 days by western blot. We found that DRD2 was specifically increased after treatment (Figure 1C), compared to the other dopamine receptors. We also performed FACS analysis of DRD2 expression in 3 different PDX cells lines over the same time course. In line with our western blot data, we observed induction of DRD2 protein expression following TMZ treatment (Figure 1D). To confirm DRD2 expression was not caused by culture conditions, we confirmed the expression of DRD2 in orthotopic xenografts from the brains of athymic nude (nu/nu) mice, as well as the presence of DRD2 mRNA in human patient samples (Extended Data Figure 1-1 Panel A). Our results corroborate several recent studies. First, GBM tumors in patients express DRD2; these levels are often elevated compared to healthy brain(Li et al. 2014). Critically, DRD2 is known to interact with EGFR signaling, a pathway frequently mutated or hyperactive in GBM(Schunemann et al. 2010; Winkler et al. 2004; Giannini et al. 2005; Mukherjee et al. 2009; Snuderl et al. 2011; Szerlip et al. 2012; Verhaak et al. 2010). Recent studies have shown that DRD2 inhibition in conjunction with EGFR antagonists reduced GBM proliferation(O’Keeffe et al. 2009; Li et al. 2014). Based on these results, we selected DRD2 as our primary receptor for further exploration.

### CD133^+^ glioma initiating cells respond to DRD2 activation with electrophysiological changes

We next tested if DRD2 expression correlates with certain cellular phenotypes. First, we examined how DRD2 protein expression is affected by alterations in culture conditions known to promote the GIC state. We examined expression of DRD2 in PDX cells growing either in 1% fetal bovine serum (FBS) containing differentiation condition media or as spheres in supplemented neurobasal media. We found that cells growing as spheres in GIC maintenance media had elevated expression not only of GIC markers like CD133 and Sox2, but also heightened DRD2 (Figure 2A), indicating that cellular state can influence this receptor expression. We next analyzed if DRD2 expression correlates with the expression of CD133, a widely used GIC marker. FACS analysis revealed that DRD2 expression is elevated in the CD133^+^ population (Figure 2B).

**Figure 2:**
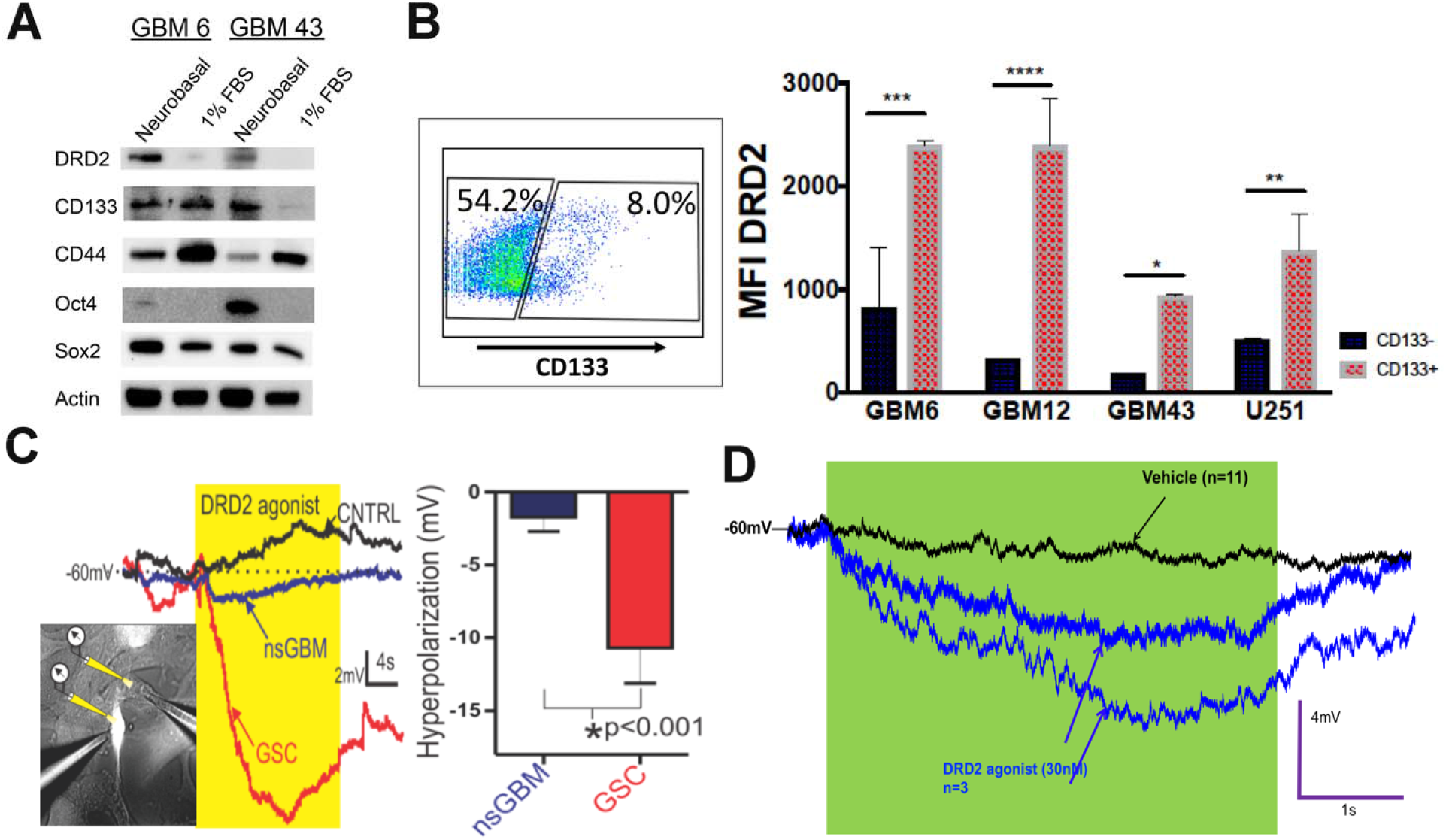
Glioma initiating cells preferentially express DRD2 and specifically respond to receptor activation with hyperpolarization. (A) Two different subtypes of PDX lines freshly isolated from animals were cultured in the tumor sphere maintenance media (neurobasal media containing EGF and FGF) or differentiation media (1% fetal bovine serum) and subjected to immunoblot analysis for the expression of DRD2 and various GIC markers. (B) PDX cells were subjected to FACS analysis for GIC marker CD133 and DRD2. Using controls, we established gates for CD133^+^ and CD133^−^ populations. We then quantified the mean fluorescence intensity (MFI) for DRD2 expression in each of these populations. (C) Human glioma U251 cells expressing red fluorescent protein (RFP) under the control of the CD133 promoter were patch-clamped and electrophysiological recordings were taken during exposure to highly selective DRD2 agonist. Puffing DRD2 agonist (30nM, yellow) onto fluorescently labeled GIC (brightly labeled cell left panel, red membrane potential trace middle panel) induces a more robust hyperpolarization of the membrane potential (mV) than non-GICs exposed to the same agonist (nsGBM, non-labeled cell left panel, blue trace middle panel), or versus puffing vehicle control (n=4 each). Graph: DRD2 agonist hyperpolarizes GIC more strongly than nsGBM cells (right panel). (D) Electrophysiological recordings of PDX cells confirms that only certain cells respond to the agonist with hyperpolarization. **Bars** represent means from three independent experiments and **error bars** show the SD. Student T-tests were performed for each day separately. *P<.05, **P<.01, ***P<.001.

Intrigued by this differential expression, we analyzed functional differences in DRD2 signaling in GICs and non-GICs within a heterogeneous cell population. One of DRD2’s main effects is to alter the polarization of brain cells(Beaulieu and Gainetdinov 2011). We therefore utilized electrophysiology to determine the effects of DRD2 activation on GBM cellular polarization. We have previously developed a GIC-reporter line in which expression of red fluorescent protein (RFP) is under the control of a GIC-specific CD133-promoter. This system enables faithful identification of tumor-initiating GICs in live cells and in real time(Lee et al. 2016). We treated our high-fidelity U251 GIC reporter cell line with the highly specific DRD2 agonist (3’-Fluorobenzylspiperone maleate, Tocris) at a dose of 30nM, the estimated concentration of dopamine in cerebrospinal fluid(Floresco et al. 2003; Keefe, Zigmond, and Abercrombie 1993). Electrophysiological recordings revealed that RFP^+^CD133^+^ cells responded to DRD2 activation to a greater extent than RFP^−^ non-GICs in the same population (Figure 2C). Notably, this activation produced hyperpolarizations similar in amplitude and duration to neuronal responses(Beaulieu and Gainetdinov 2011). We also confirmed that only certain PDX cells respond to the agonist in PDX cells (Figure 2D). These results show that DRD2 receptors in GBM cells are functional and produce GIC-specific effects.

### Activation of DRD2 increases GBM cell self-renewal and tumor engraftment capacity

We next sought to understand the relationship between DRD2 expression and tumor cell phenotype, specifically self-renewal ability and tumor engraftment capacity. To that end, PDX GBM cells were treated with either DMSO or TMZ. Following four days of exposure, cells were sorted based on DRD2 expression and plated for limiting dilution neurosphere assays. We found that DRD2 positive (DRD+) cells exhibited about 4-fold elevated sphere formation capacity as compared to DRD2 negative (DRD−) cells (Figure 3A), suggesting that the DRD2+ population has enhanced self-renewal capacity. Exposure to physiological dose of TMZ also enhanced the self-renewal capacity in both DRD2+ as compared to DRD2-population. Most importantly, post TMZ therapy enhancement of self-renewal resulted in enhanced tumor engraftment capacity in the orthotopic xenograft model as 100% of the mice developed tumor when 500 DRD+ cells were implanted in the brain of the immunocompromised nu/nu mice (Figure 3B).

**Figure 3:**
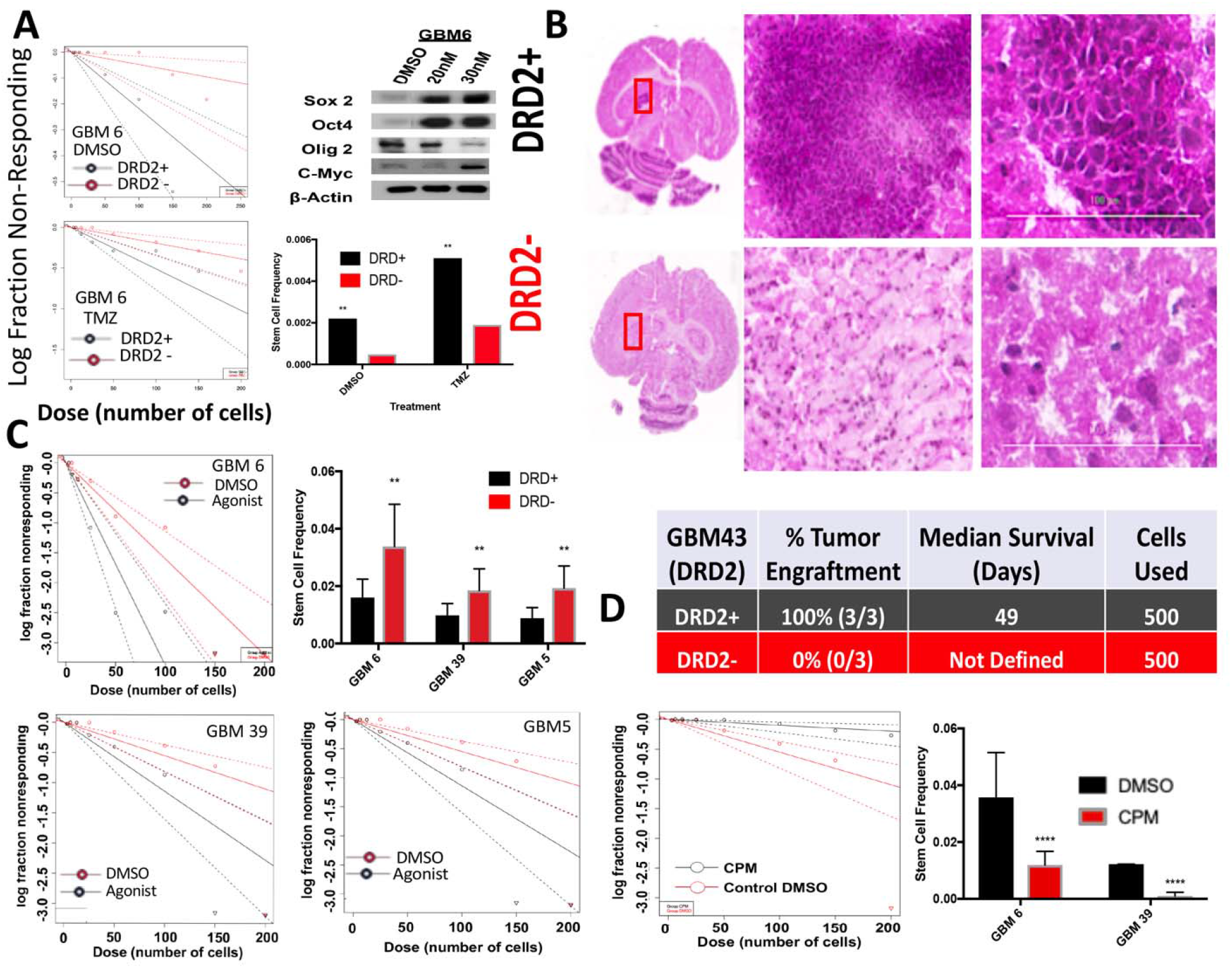
Activation of DRD2 signaling increases the self-renewal capacity of GBM cells. (A) PDX GBM cells treated with either DMSO or TMZ were sorted based on DRD2 expression and neurosphere assays were performed. Using the extreme limiting dilution assay algorithm, frequency of GICs was calculated (DMSO, DRD2+ 1/458 and DRD2− 1/2091, p<.001; TMZ, DRD+ 1/196 and DRD2− 1/515, p<.001). (B) In vivo engraftment of 500 cells sorted on DRD2 + or − expression demonstrates DRD2+ cells engraft more efficiently. Extended data figure 3-1 includes more images of engrafted tumors from other mice. PDX lines treated with dopamine agonist demonstrate elevation in key stem cell genes via western blot. Extended data figure 3-1 includes western blots from other PDX lines demonstrating similar trends. (C) PDX GBM cells were treated with either DRD2 agonist (30nM) or equimolar vehicle control DMSO and plated for neurosphere assays. Using the extreme limiting dilution assay algorithm, frequency of GICs was calculated (GBM6, DMSO stem cells 1/62 and agonist stem cells 1/29.2 (p<.01); GBM39, DMSO stem cells 1/181.9 and agonist stem cells 1/87.8 (p<.05); GBM5, DMSO stem cells 1/118.9 and agonist stem cells 1/67.4 (p<.05). Extended data figure 3-1 includes a non-responding GBM12 neurosphere assay. (D) Two PDX lines were treated with either chlorpromazine (CPM, 1 μM) or equimolar DMSO and plated for neurosphere assays as above. Difference between groups was determined by score test of heterogeneity. Error bars represent upper and lower limit of 95% confidence interval computed for stem cell frequency. Student t-Tests were performed for each comparison. ****P<0.0001.

Based on our data described above as well as recently reported data demonstrating that DRD2 signaling influences normal progenitor phenotype and differentiation(Baker, Baker, and Hagg 2004; O’Keeffe et al. 2009; Winner et al. 2009; Ohtani et al. 2003), we next examined if the observed self-renewal capacity of DRD2+ cells was indeed DRD2 dependent and if subtypes influence such characteristic. Several different subtypes of PDX cell lines were plated in the presence of either vehicle control DMSO or 30nM DRD2 agonist and, after 8 days, neurospheres were counted. Our results demonstrate that, for three of the PDX lines tested, DRD2 activation increases sphere-forming capacity (Figure 3C). GBM12, however, did not respond to the agonist, thus indicating may be some subtype specific variability when responding to DRD2 signaling (Extended Data Figure 3-1) In order to confirm this increase in GIC, we analyzed the expression of several key GIC markers following treatment with the DRD2 agonist and found that cell lines that responded to the agonist by increasing sphere formation also showed increased expression of genes associated with stemness (Extended Data Figure 3-1). Importantly, treatment did not increase cell proliferation as measured by the Ki67 FACS, suggesting our result comes from cellular plasticity and not increased GIC proliferation (Extended Data figure 3-1). Additionally, inhibition of DRD2 by chlorpromazine (CPM, 1μM), a selective DRD2 antagonist(Beaulieu and Gainetdinov 2011) in agonist-responsive cell lines GBM6 and GBM39, markedly reduced sphere formation, as measured by neurosphere assays (Figure 3C).

### Glioblastoma cells express tyrosine hydroxylase (TH) and secrete dopamine

Our data indicate that DRD2 activation alters the phenotype of GBM cells. We next investigated the possibility that this DRD2 activation occurs via autocrine release of its natural ligand, dopamine. Production of monoamines like dopamine is usually a tightly regulated process limited to specific dopaminergic neuronal populations. Critically, all our experiments were performed in purified monocultures of cancer cells in the absence of dopaminergic neurons, raising the question of the mechanism of DRD2 activation. To investigate the possibility that tumor cells synthesize their own dopamine, we first examined expression of tyrosine hydroxylase (TH), the rate-limiting enzyme of dopamine synthesis, in PDX GBM cells (Figure 4A). To confirm this expression occurs *in vivo*, we intracranially implanted PDX GBM cells into athymic nude mice; upon visible signs of tumor burden, mice were sacrificed and whole brains were obtained. FACS analysis revealed that human leukocyte antigen (HLA+) tumor cells do in fact express TH *in vivo* (Figure 4B). Examination of GlioVis data(Bowman et al. 2017) also demonstrated that patient GBM samples express TH and other enzymes necessary for dopamine synthesis, as well as transporters for vesicle loading and reuptake (Extended Data Figure 4-1).

**Figure 4:**
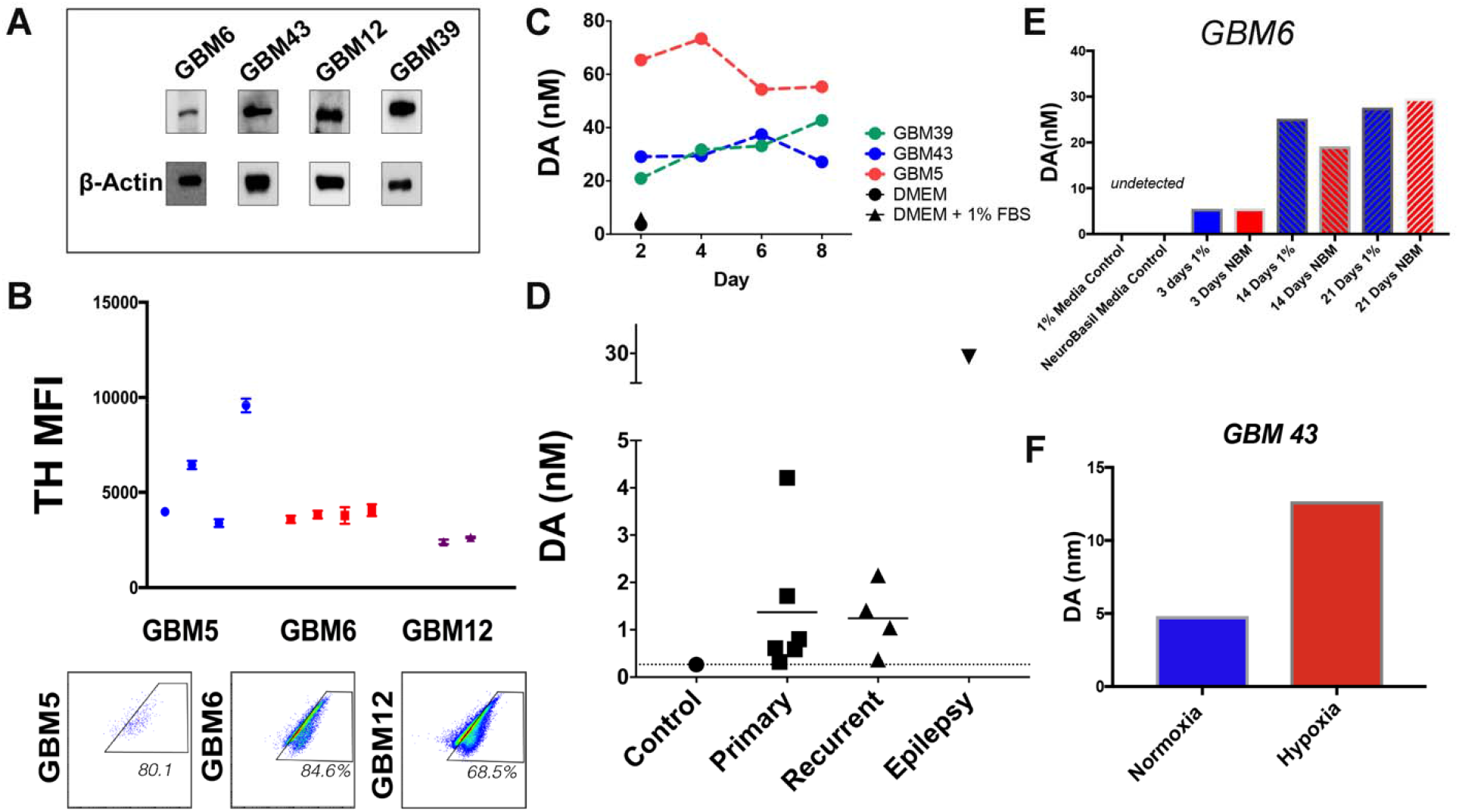
GBM cells synthesize and secrete dopamine. (A) Western blot analysis of tyrosine hydroxylase (TH) expression, the rate-limiting enzyme in dopamine synthesis. Beta-actin was used as a loading control. (B) PDX cells were implanted intracranially into athymic nude mice. Upon sickness from tumor burden, mice were sacrificed and whole brains extracted. FACS analysis of human leukocyte antigen positive tumor cells demonstrates that GBM tumors continue to express TH in vivo. Each dot represents individual mice; FACS were run in technical triplicates. In set scatter plots are representative for FACS results in each cell line tested. (C) Conditioned media from PDX GBM cells growing in purified monocultures were analyzed by high performance liquid chromatography with mass spectrometry (HPLC-MS) for dopamine. Unconditioned DMEM media with and without 1% FBS was used as a control. (D) HPLC-MS was performed on GBM samples from patients undergoing resection. Cortical tissue from an epilepsy surgery was used as the control. (E) PDX GBM6 cells were growing in either 1% FBS containing media or Neurobasal media (NBM). Conditioned media were collected at various days and dopamine levels were calculated using HPLC. (F) PDX GBM43 cells were cultured in the presence of either 20% or 1% oxygen (hypoxia) for three hours. Conditioned media was collected, and dopamine levels determined. **Dots** represent values from each timepoint and/or sample tested. Extended data figure 4-1 includes other genes involved in dopamine synthesis (VMAT2, DAT, DDC, TH) measured in patient tumor samples as well as HPLC preformed on various PDX subtypes undergoing a TMZ treatment time course.

To determine if PDX monocultures were generating dopamine, we assayed dopamine levels using high performance liquid chromatography with mass spectrometry (HPLC-MS). Conditioned media of PDX monocultures were found to have dopamine levels of 20-65 nM, between 10 and 30 times greater than cell free media (Figure 4C). The levels of dopamine were relatively stable in samples taken over 8 days, suggesting that the PDX cells produce and maintain basal dopamine synthesis. Next, we quantified dopamine in human tumor samples. A portion of cortex removed from an epilepsy surgery were also analyzed to determine dopamine levels in non-cancerous brain tissue. These assays revealed dopamine expression in all cases (Figure 4D).

In light of our previous data indicating that dopamine signaling is associated with cell state, we next examined how cell state influences dopamine secretion. PDX GBM6 cells were cultured in either 1%FBS or neurobasal media. Conditioned media were collected after 3, 14, and 21 days and dopamine levels determined by HPLC-MS. We found that dopamine production was time dependent, but not clearly linked to culture conditions (Figure 4E). It has been established that hypoxia induction can influence the ability of stem cells to differentiate and produce dopamine(Wang et al. 2013). We therefore analyzed dopamine levels from media in which PDX GBM cells were growing exposed to either 20% normoxia or 1% hypoxia. HPLC-MS revealed that hypoxia increase dopamine levels (Figure 4F), suggesting a similar mechanism.

### Activation of DRD2 induces expression of HIF proteins

To determine how DRD2 signaling may underlie the observed changes in self-renewing capacity, we analyzed patient tumor samples from The Cancer Genome Atlas (TCGA). Patient samples were stratified from lowest to highest expression of DRD2 and correlations were determined for 12,042 genes by Pearson correlation coefficient analysis; coefficients >0.5 or <0.5 and false discovery rate (FDR) <0.05 were used. Genes fitting these parameters were then analyzed using Enrichr pathway analysis program. Our analysis revealed that DRD2 levels are positively correlated with the expression of hypoxia inducible factors (HIF) 1α and HIF2α pathways, as well as with the Reelin and ErbB4 signaling networks (Figure 5A).

**Figure 5:**
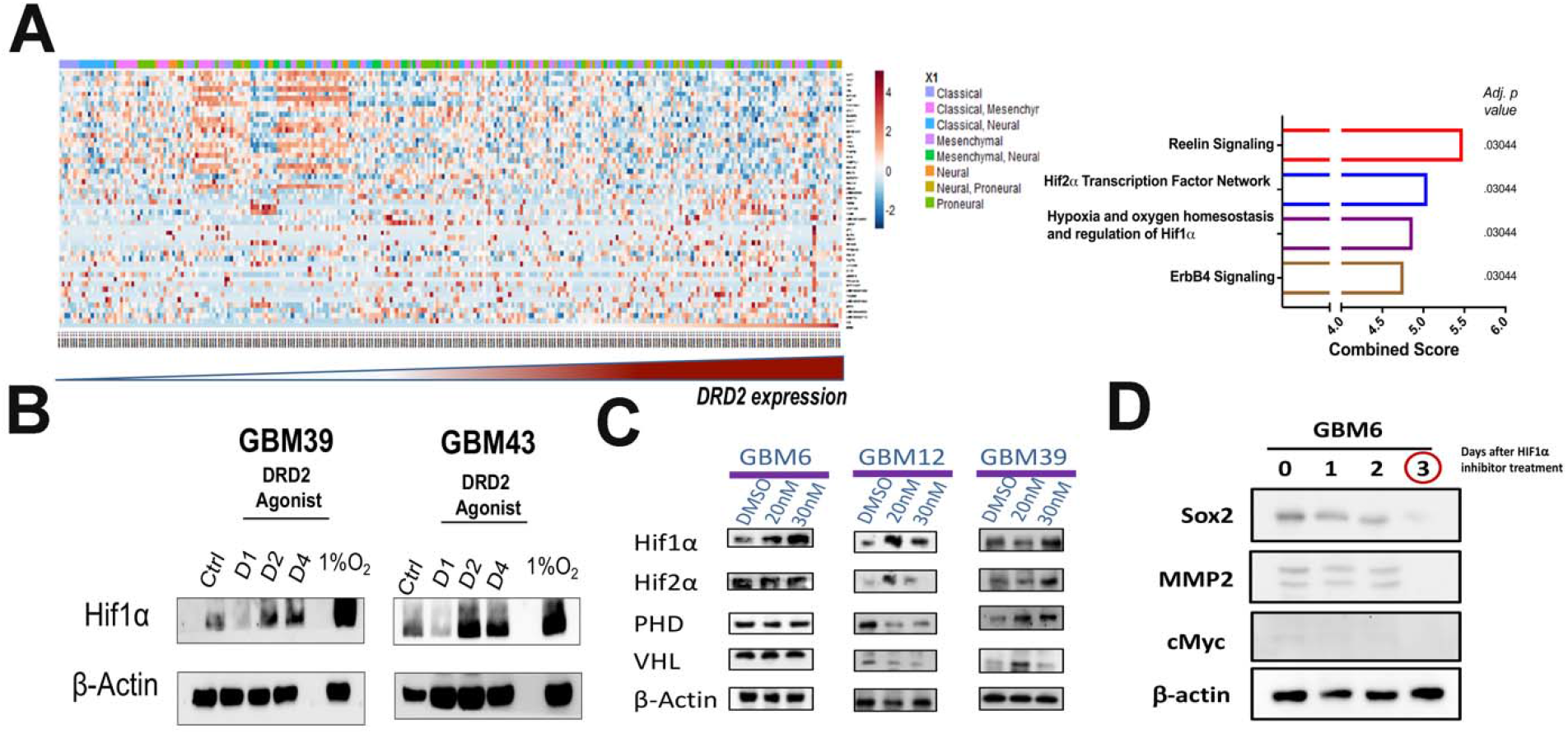
DRD2 signaling activates hypoxia inducible factors in normoxic cultures. (A) Correlation between DRD2 and 12042 genes from TCGA was determined by Pearson correlation coefficients. 49 genes with coefficients >0.5 or <-0.5 and false discovery rate (FDR) <0.05 were selected. Genes fitting these parameters were then analyzed using Enrichr. Top hits are shown in the graph, with combined score and adjusted p value. (B) PDX GBM cells were treated with DRD2 agonist (30nM) for 1, 2, or 4 days and then probed for expression of HIF1α by immunoblot analysis. DMSO treated cells as vehicle treated control and cells exposed to hypoxic conditions (1.5% O2) were used as a positive control for HIF expression, and beta actin was used as a loading control. (C) Western blots were used to analyze the level of proteins in the VHL-PHD regulatory pathway after treatment with either DMSO or DRD2 agonist (20nM or 30nM). Beta actin was used as a loading control. (D) PDX GBM6 were exposed to DRD2 agonist (30nm) and treated with HIF inhibitor for 24 48 and 72 hours SOX2, MMP2, and CMYC protein levels were assayed.

Hypoxia is well known to influence the phenotype of GBM cells, pushing them towards a more GIC phenotype(Evans et al. 2004; Soeda et al. 2009). Further, we have previously shown that GBM cells utilize HIF signaling to promote cellular plasticity following therapy(Lee et al. 2016). To investigate how DRD2 stimulation affects HIF signaling, PDX cells GBM39 and GBM43 were treated with agonist under normal and hypoxic conditions. Immunoblot analysis demonstrated a substantial, time-dependent increase in HIF1α levels in both cell lines following activation of DRD2 signaling, regardless of oxygen availability (Figure 5B).

HIF protein levels are controlled both at the level of gene transcription and via hydroxylase-dependent degradation. In order to determine if DRD2’s activation of HIF was occurring at the level of gene transcription or degradation, we analyzed the levels of PHD and VHL, key regulators of HIF stabilization, following treatment with the DRD2 agonists by western blot. These assays revealed cell line specific changes in the HIF pathway. In GBM12, the expected correlations were observed—elevated HIF levels matched decreased levels of both PHD and VHL. However, in GBM39, the increased HIF expression actually correlated with elevated PHD and VHL (Figure 5C). Notably, these results indicate that increases in HIF levels following DRD2 activation occurs in both cell lines that are responsive and non-responsive in neurosphere assays. This common shared HIF induction with disparate functional outcomes suggests that shared activation of HIF leads to unique downstream effects.

We next sought to determine if the increased GIC population following DRD2 activation was dependent on the previously identified increase in HIF1α in the clinically relevant PDX model. GBM 6 treated with HIF1α inhibitor after exposure to DRD2 agonist show loss of Sox2, CMYC, and MMP2 expression (Figure 5D).

### Activation of DRD2 induces alterations in expression of HIF-responsive genes and functional changes in metabolic phenotype

HIF signaling can have profound effects on cancer cells, ranging from shifts in gene expression to altered functional attributes. For a full transcriptome profile of the mechanisms governing DRD2’s influence on GBM cells, we performed microarray analysis of gene expression in cells treated with either vehicle control DMSO or 30nM DRD2 agonist for four days. We chose two cell lines with distinct responses to dopamine in sphere-forming assays—neurosphere-responsive GBM39 and non-responsive GBM12. To determine the changes that were specific to each cell line, we utilized the online GeneVenn tool. Transcripts with a p value <.05 from each cell line, comparing agonist and DMSO controls, were identified and overlap was analyzed by GeneVenn. We found that 3479 increases in gene expression were specific to neurosphere-responsive GBM39, while 1917 increases were unique to neurosphere-non-responsive GBM12. These two cell lines showed shared increases in only 287 genes (Figure 6A). Enrichr analysis of these cell lines’ dopamine-induced transcriptomes revealed activation of unique signaling pathways in each cell line (Figure 6B). Given our result indicating a connection between HIF signaling and DRD2, we next examined genes known to be regulated by HIF during canonical hypoxia responses. We therefore performed GSEA analysis of only the hypoxia response gene-sets. This analysis confirmed a subtype-dependent response to DRD2 activation. GBM39 cells did not activate canonical HIF targets following agonist, whereas GBM12 did (Figure 6B). These results indicate a specific change in gene expression following the agonist and suggest that these differences may underlie the functional changes in sphere-formation.

**Figure 6:**
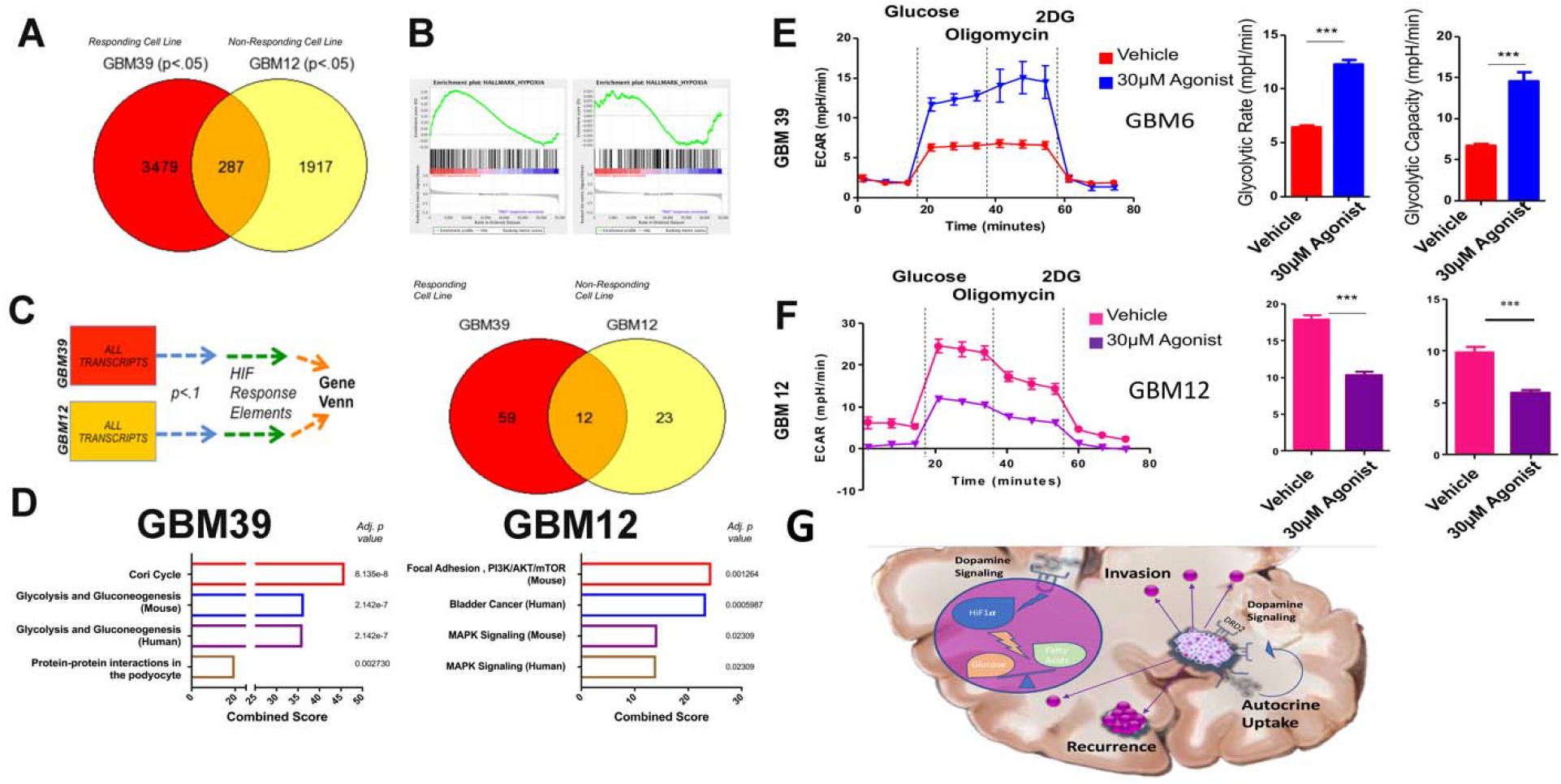
DRD2 triggers alterations in gene expression and metabolic phenotype. (A) Microarray analysis of gene expression was performed in PDX cells treated with DRD2 agonist after four days. GeneVenn analysis was performed on gene transcripts found to be significantly elevated in the two cell lines, revealing subtype specific alterations in expression profiles. (B) Gene set enrichment analysis (GSEA) was performed on each cell line for canonical hypoxia response signaling. (C/D) Microarray results were then limited to genes with confirmed hypoxia response elements. (E/F) Seahorse analysis was performed to quantify the glycolytic rate in two PDX lines, (E) GBM39, which responds to DRD2 activation with increased sphere-formation, and (F) GBM12, which are unresponsive to DRD2 activation in neurosphere formation. Cells were treated with either DMSO or 30nM DRD2 agonist and subjected to Seahorse analysis after 96 hours. Extended data figure 6-1 includes a responding PDX line, GBM5, as an assay control demonstrating similar effects. Figure 6-1 also includes FACS analysis of glucose uptake and fatty acid uptake in GBM12 and 39 as further validation of seahorse assays. (G) Summary schematic of dopamine’s role within the tumor microenvironment and at the single cell level. **Microarray** was performed on biological triplicates. Bars represent means from three independent experiments and error **bars** represent SD. Student T-tests were performed for each separate cell line. *P<.05, **P<.01, ***P<.001.

While many genes are activated in response to HIF proteins, certain genes are known to be activated via direct binding of HIFs to gene promoters at conserved regions termed HIF-responsive elements (HRE). To determine if the observed changes in gene expression occurred at the level of actual HIF binding, we collected all the gene transcripts that were significantly upregulated after treatment with the DRD2 agonist and known to have a promoter containing the HRE (previously confirmed by ChIP-Seq(Schodel et al. 2011)). These sets of genes were then analyzed using the Enrichr platform. Once again, we found that the two cell lines showed drastically different responses to the agonist (Figure 6C). Treated GBM39 cells showed increased levels of HIF-regulated genes, including glycolysis, whereas GBM12 showed increased activation of the PI3K/AKT pathway (Figure 6D). These results suggest that the differential effects of DRD2-induced activation of HIF in classical GBM39 and proneural GBM12 depend on unique signaling mechanisms.

Changes in gene expression, while informative does not guarantee a functional alteration in cellular behavior. We therefore sought to determine if activation of DRD2 does in fact lead to GBM39-specific alterations in metabolism. First, we analyzed the uptake of glucose and fatty acid via FACS, utilizing 2-NBDG, a well-established fluorescent analog of glucose, and our proprietary fatty-acid tagged quantum dots(Muroski et al. 2017). Only GBM39 showed altered fatty acid uptake following DRD2 activation. However, all cells treated with the agonist showed increases in glucose uptake (Extended Data Figure 6-1). To further characterize the effects of DRD2 signaling on glucose metabolism, we performed extracellular flux analysis. We discovered a differential effect predicted by each cell line’s behavior in neurosphere assays. GBM39, of the classical subtype and responsive to the agonist in neurosphere assays, showed a marked increase in glycolytic rate as measured by glucose stimulated extracellular acidification (ECAR) (Figure 6E). In contrast, non-responsive and pro-neural subtypes GBM12 and GBM43 actually showed decreases in glycolysis following DRD2 agonist treatment (Figure 6F).

## Discussion

In this study, we investigated how DRD2 activation contributes to the molecular and functional phenotype of GBM. First, we demonstrated that therapeutic stress induced by anti-glioma chemotherapy alters the epigenetic status of the DRD2 promoter and subsequently increases DRD2 protein expression. Notably, this expression was elevated in GICs and could be induced by GIC-promoting culture conditions. In classical GBM cells, DRD2 activation increased sphere forming capacity, an indicator of GIC state and tumorgenicity, as well as the expression of several key GIC markers. We further found that GBM cells themselves can generate dopamine in purified monocultures, and therefore exhibit autocrine/paracrine DRD2 activation capabilities. In all lines tested, activation of DRD2 induced the expression of HIF proteins, regardless of oxygen availability. Whereas the HIF induction by DRD2 activation was common to all cell lines tested, the downstream functional changes varied. Classical GBM39 cells exhibited increased uptake of glucose and glycolytic rate following DRD2 stimulation. In contrast, proneural GBM12 cells showed no change in sphere-forming capacity. However, activation of DRD2 did induce expression of canonical HIF-responsive genes and led to a reduction in their glycolytic rate. Cells exposed to low oxygen conditions typically activate HIF and subsequently upregulate glycolysis; the fact that proneural cells do not upregulate this metabolism, despite elevated HIF levels, suggests some unique HIF regulatory mechanism. In sum, these results indicate a key role for DRD2 signaling in influencing GBM cellular phenotype, at the level of both gene expression and functional attributes.

Induction of cellular plasticity, specifically in steering cells towards a treatment-resistant GIC state, is a key problem in GBM and contributes to tumor recurrence(Safa et al. 2015; Olmez et al. 2015; Dahan et al. 2014; Lee et al. 2016). Our data demonstrate that dopamine signaling plays a role in this cellular conversion. Functionally, exposure to DRD2 agonist was sufficient to increase sphere-forming capacity of PDX GBM cells. Given the concurrent increase in GIC marker expression, this treatment appears to induce the conversion of GBM cells to a more stemlike state.

Our results indicate that activation of DRD2 induces unique downstream effects dependent on the genetic subtype of cell line used. Even more interestingly, these unique changes in gene expression and phenotypes occurred despite shared activation of a key signaling node—hypoxia inducible factors. GBM39, of the classical tumor subtype, responds to the agonist with increased sphere-formation, expression of key GIC markers, altered gene expression related to metabolism, and increased glycolytic rate. In contrast, GBM12, a proneural cell line, did not increase neurosphere formation or expression of GIC markers, but did show induction of HIFs and altered gene expression related to canonical HIF signaling. It remains an open question as to which other signaling mechanisms when activated in these two cell lines enable common signaling nodes to induce cell-line specific changes in gene expression and functional phenotype. The unique genetic makeup of each cell line may provide an explanation of this phenomenon. Analysis of patient sample gene expression data and orthotopic xenograft tumor protein levels showed that proneural tumors and GBM12 xenografts have somewhat elevated expression of DRD2 (Extended Data Figure 1-1), suggesting that this signaling axis may already be active. Further, GBM39 is of the classical subtype and carries the EGFRvIII mutation, while GBM12 expresses wildtype EGFR(Giannini et al. 2005). EGFR is connected to dopamine signaling in neural development(Hoglinger et al. 2004); therefore, the fact that EGFR mutant cell lines respond with increased sphere-formation suggests that increased EGFR activity may play a critical role in the ability of DRD2 to alter cellular plasticity.

These differential responses to activation of the same receptor further highlight a key challenge facing the neuro-oncology community—intratumoral heterogeneity. GBM tumors have been shown to contain cells corresponding to each of the three subtypes within a single tumor(Sottoriva et al. 2013; Patel et al. 2014). This wide range of cell types and diverse molecular phenotypes currently limit the effective treatment of GBM. These data highlight the importance of this heterogeneity, by suggesting a mechanism for the same molecular signal to exert unique influences on different subpopulations of tumor cells. It is not hard to imagine that dopamine may generate multiple adaptive features, including elevated self-renewal capacity or maintenance of the GIC-pool via increased glycolysis, and neo-angiogenesis in GBM cells.

These data illustrate that dopamine signaling exerts influence on global as well as HIF-dependent gene expression. Further, our data demonstrate that these changes in gene transcription have a functional effect, as dopamine signaling is capable of altering the metabolism of GBM cells. GBM39 showed a highly elevated glycolytic rate following agonist treatment, while GBM12 cells had slightly reduced rates of glycolysis. How this increase in glucose uptake and glycolytic rate relates to induction of the GIC state, however, remains an open question. Several groups have recently demonstrated that altered metabolic phenotypes can influence the acquisition of a stem cell phenotype in cancer cells(Currie et al. 2013; Mao et al. 2013); it is not outside the realm of possibility that a similar effect is at play here. Further research will delineate the precise connection between elevated glucose uptake and glycolytic rate and altered cell state.

It should be noted that a recent report suggests that dopamine may actually inhibit growth of glioma. Lan and colleagues showed that dopamine reduced growth of U87 and U251 cells growing as subcutaneous xenografts(Lan et al. 2017). While difference in cell lines (U87 v. PDX) as well as the subcutaneous placement of tumors may account for this difference, this result suggests a dose-dependent effect of dopamine. Critically, our experiments were performed with doses of DRD2 agonist between 20 and 30nM, while Lan *et al* performed experiments with doses of dopamine between 10-25. Dopamine receptor signaling is highly sensitive to concentration of ligands and antagonists, which complicates understanding its role in GBM (reviewed(Caragher et al. 2017)). It is possible that the influence of dopamine signaling is dose-dependent. Further, this work highlights the complex nature of dopamine’s multiple receptors. Adding a pan dopamine receptor agonist, as they did, will activate all five receptors, which have unique downstream signaling pathways.

Our data also show that GBM cells can synthesize and secrete dopamine, an ability assumed to be the sole province of neurons. However, there is mounting evidence that GBM cells have the ability to adopt certain neuronal functions. Osswald et al. demonstrated that GBM cells utilize GAP43, neuronal outgrowth cone protein, to form networks(Osswald et al. 2015). Further, it has been shown that GBM cells synthesize and secrete glutamate, an excitatory neurotransmitter, and alter cortical activity(Buckingham et al. 2011). Developmental neurobiology also provides some clues as to how these cells can attain a neuron-specific behavior. Neural stem cells engrafted into the striatum of mice can spontaneously attain dopaminergic traits, suggesting that secreted factors in the brain can induce the formation of dopamine secreting cells(Yang et al. 2002). Given the highly similar expression profile of NSCs and GSCs, a similar mechanism may induce the formation of a dopamine-producing population of glioma cells. Finally, a recent report utilizing TCGA datasets indicated that high TH expression correlates with reduced median survival(Dolma et al. 2016). Critically, this data analyzed mRNA expression in GBM cells themselves, not in surrounding brain cells. Therefore, the survival difference relies on how TH functions in the tumor.

In short, GBM cells exhibit a variety of neuron-like properties, including the ability to synthesize and secrete dopamine. Our *in vitro* data indicate that the GBM cells are synthesizing dopamine rather than taking up dopamine from the surrounding dopamine neurons and storing it. Our results clearly fit within this emerging theory that GBM tumors may display neuron-specific behaviors. More research is needed to delineate the specific subpopulations of GBM tumors that secrete dopamine and the mechanisms governing this process. This result, combined with the evidence that DRD2 signaling promotes a shift to the GIC state, suggests that tumors utilize dopamine as a mechanism to control their microenvironment and promote a pro-growth, pro-stemness milieu.

It could be argued that even if GBM cells do utilize dopamine in an autocrine manner, they must also rely on ambient dopamine in the brain microenvironment to induce major signaling changes. It remains possible that non-tumor dopamine also influences GBM. It has already been shown that proteins from the synaptic cleft of nearby neurons can influence glioma growth(Venkatesh et al. 2015). We therefore cannot rule out the possibility that our tumor sample data represent cells that have absorbed dopamine rather than produced it (Figure 4). Indeed, TCGA indicates that GBM cells express the dopamine transporter (Extended Data Figure 4-1). In addition, it has been shown that patients with Parkinson’s Disease, characterized by the loss of dopaminergic neurons, have a lower incidence of high grade gliomas(Diamandis et al. 2009). Further, studies of tumor localization indicate that GBM tumors preferentially localize to areas of dopamine innervation(Larjavaara et al. 2007). We do not exclude or reject the possibility that ambient dopamine influences tumor behavior, with two key caveats. First, given that only specific portions of the brain contain dopaminergic neurons(Ungerstedt 1971) and that GBM tumor growth often disrupts the normal neural circuitry in their area, it cannot be guaranteed that dopamine from the microenvironment is widely available for every tumor. Second, the threedimensional shape of the tumor is likely to limit perfusion of certain molecules from distal sources into the cleft. As such, ambient dopamine released from neurons may also influence signaling in GBM tumors.

Finally, it remains a lingering question the extent to which the effects of dopamine signaling observed here are GBM specific or represent a recapitulation of the behavior of normal neural stem cells (NSCs) and brain progenitor cells. It has been demonstrated, for example, that NSCs in the subventricular zone respond to dopamine with increased proliferation(Baker, Baker, and Hagg 2004; O’Keeffe et al. 2009; Winner et al. 2009). Activation of dopamine receptors on progenitors in the developing brain has also been shown to expand neuronal populations and neurite outgrowth^(Yoon and Baik 2013)^, and alter cell cycle status(Ohtani et al. 2003). Finally, dopamine signaling in oligodendrocyte precursors (OPCs) has been shown to expand the pluripotent population(Kimoto et al. 2011). These studies suggest that GBM cells have hijacked an existing role for dopamine in brain development for their own gains. The connection of metabolism, however, was not identified in these studies or other relevant literature. Further research will undoubtedly shed light on this interesting connection.

In sum, our data indicate that GBM cells respond to DRD2 activation at the level of both gene expression and functional phenotype. Further research will shed light on the dynamics of tumor-derived dopamine and mechanisms influencing key factors like rate of production and secretion, interaction with non-cancerous cells in the environment, and the effects of tumor-derived dopamine on therapy resistance. Overall, this study provides new insight into how GBM cultivates a supportive microenvironmental niche to colonize the CNS and suggests that GBM tumors possess a remarkable ability to hijack cellular mechanisms previously thought to be the exclusive provenance of neurons. Given the growing chorus of studies indicating that dopamine antagonists inhibit GBM growth(Kang et al. 2017; Karpel-Massler et al. 2015; Shin et al. 2006), these results provide critical context for the source of tumor-influencing dopamine and the mechanism underlying dopamine’s role in gliomagenesis.

---

**Extended Data Figure 1-1:**
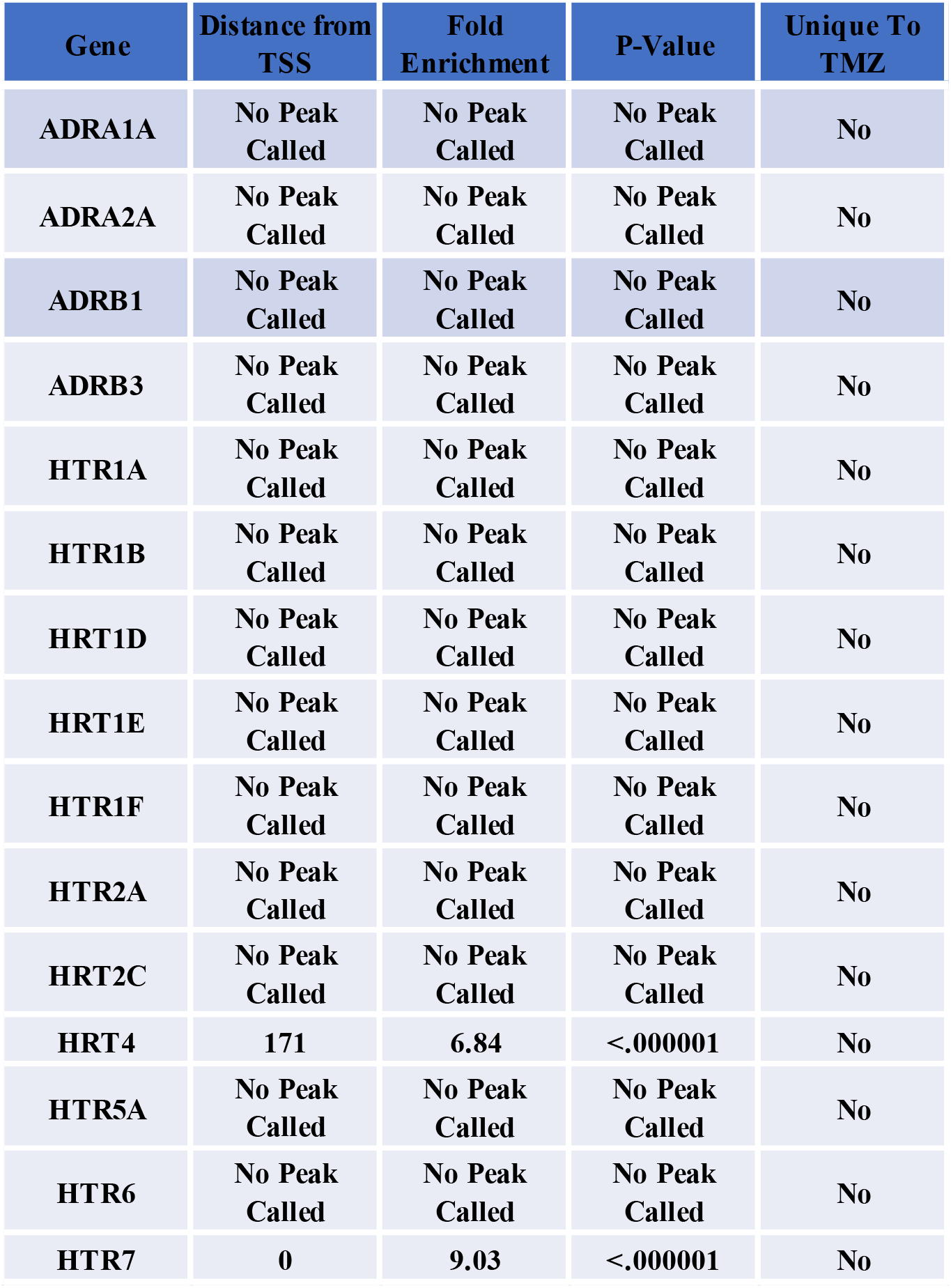
ChIP analysis of H3K27ac and H3K27me for other monoamine receptors.

**Extended Data Figure 1-2:**
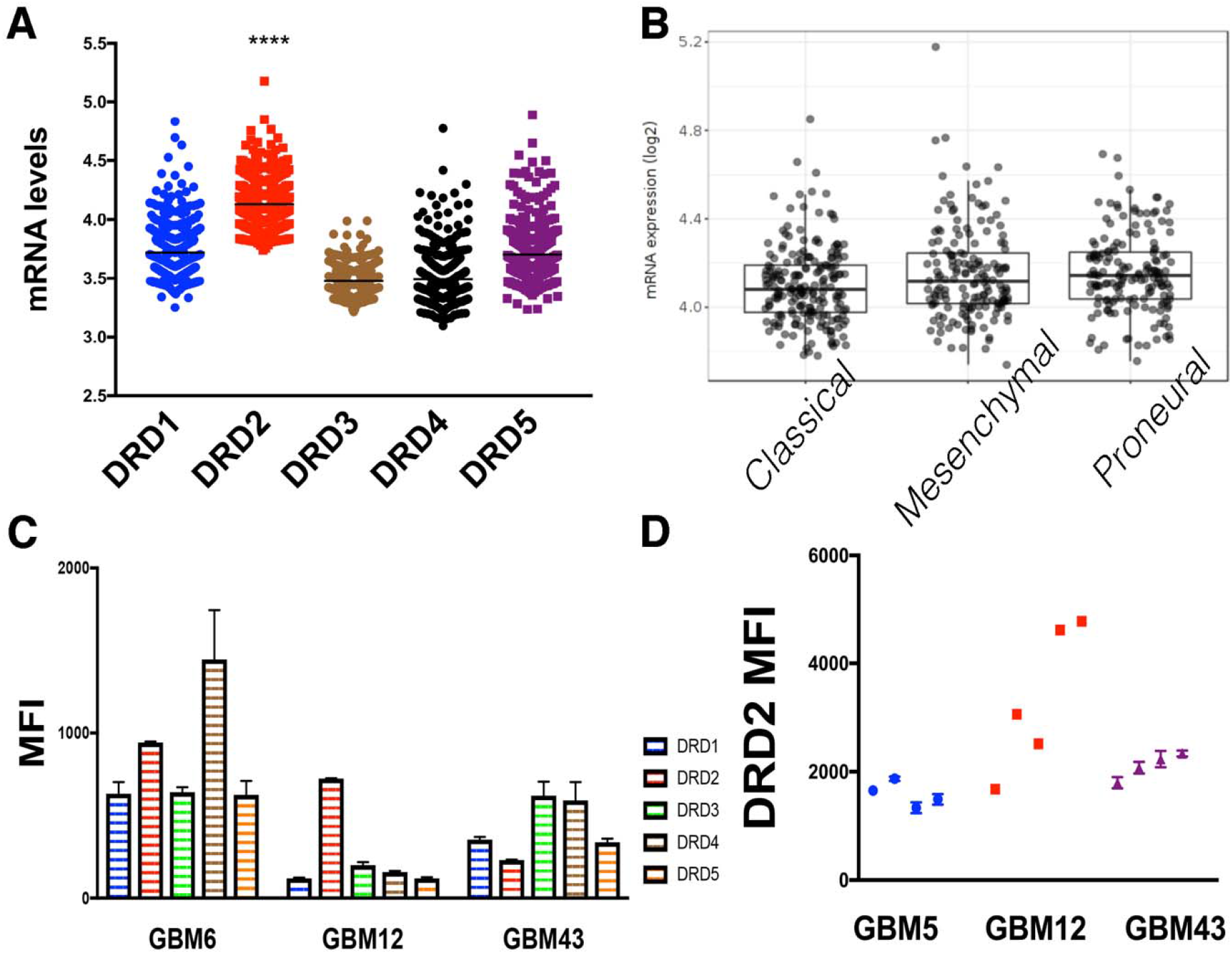
GBM tumors express DRDs. (A) TCGA data reveals elevated expression of DRD2 relative to the other DRDs. Dots represent mRNA levels from individual tumors. Mean levels were compared using one-way ANOVA. (B) DRD2 expression in patient samples varies based on subtype. (C) Expression of DRDs in PDX GBM cells was determined using FACS analysis. Bars represent the mean of three individual experiments. (D) PDX GBM cells were implanted intracranially into the right hemisphere of athymic nude mice. Dots represent the mean from each individual mouse.

**Extended Data Figure 3-1:**
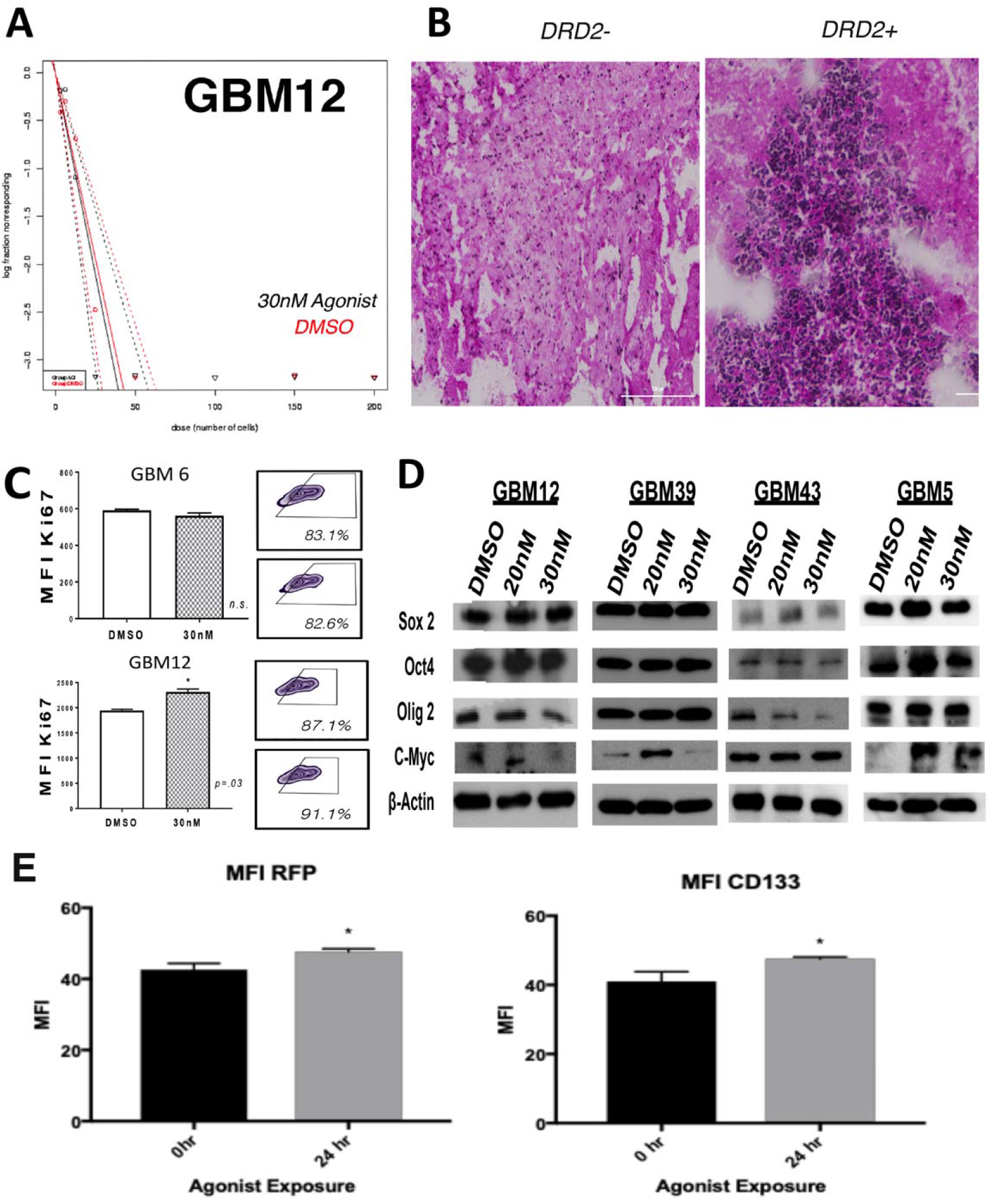
Activation of DRD2 increases expression of GIC markers. (A) Neurosphere assay of PDX GBM12 treated with either DRD2 against (30nM) or equimolar DMSO. Extreme limiting dilution assays reveal no significant difference in stem cell frequency. (B) Mice injected with DRD2+ cells show efficient tumor engraftment compared to DRD2-cells. (C) PDX GBM cells treated with agonist or DMSO and Ki67 levels were determined by FACS. No difference in Ki67 was detected. (D) PDX cells treated with agonist or DMSO and the expression of GIC markers was examined by western blot after four days. Beta-actin was used as a loading control. (E) U251-CD133 RFP driven reporter line was treated with agonist for 24 hours and RFP (reporter) and CD133 (ligand) were assayed.

**Extended Data Figure 4-1:**
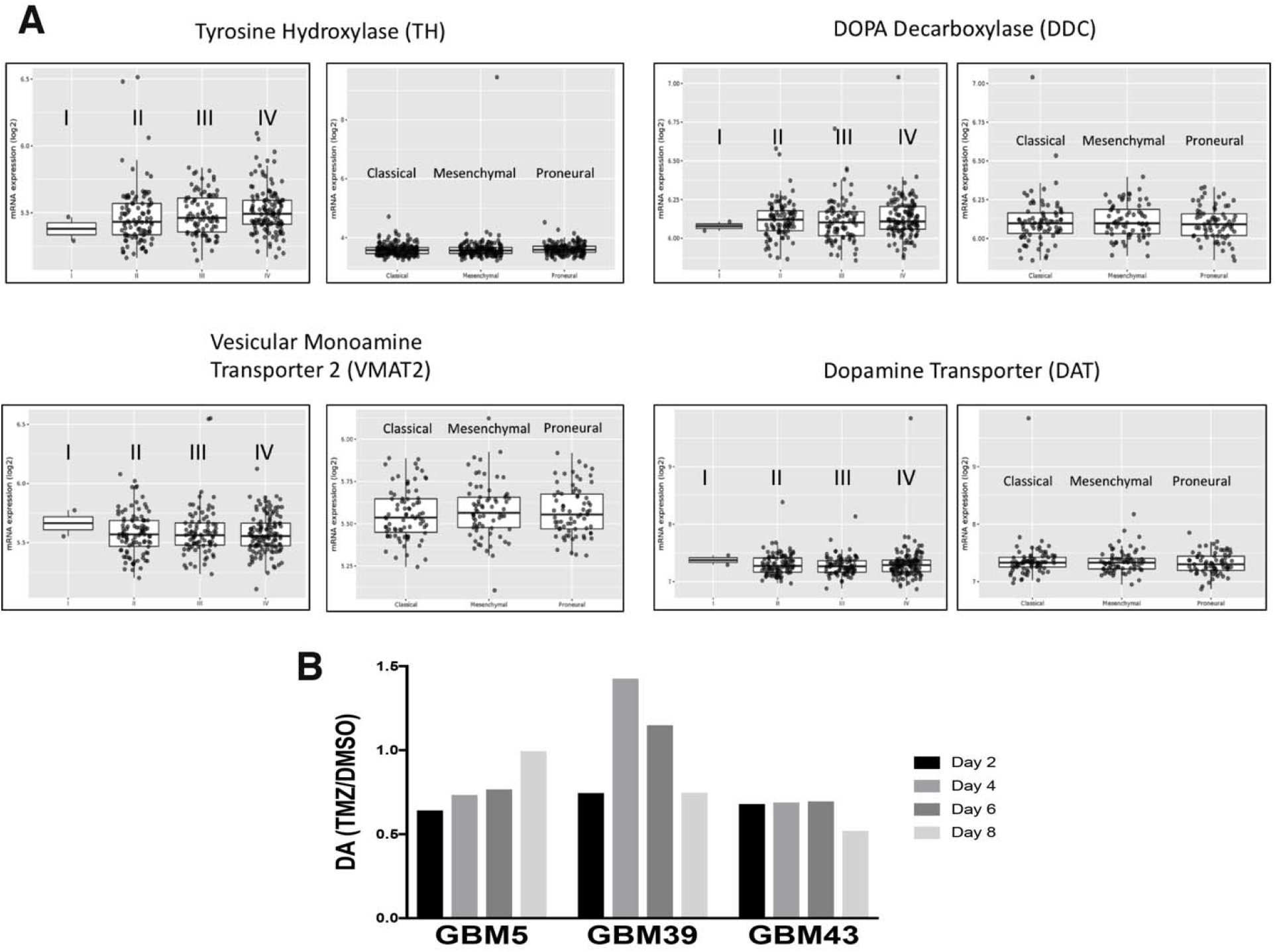
(A) Patient tumors express mRNA for several genes involved in synthesis, secretion, and reuptake of dopamine. Data were taken from the Cancer Genome Atlas (TCGA). Expression was examined in astrocytomas (Grade 1, II, III, IV) and within GBM based on subtype (Classical, Mesenchymal, Proneural). (B) PDX lines were exposed to 50μm TMZ and collected after 2, 4, 6, and 8 days and subjected to HPLC. Results are shown normalized to equimolar DMSO controls.

**Extended Data Figure 6-1.**
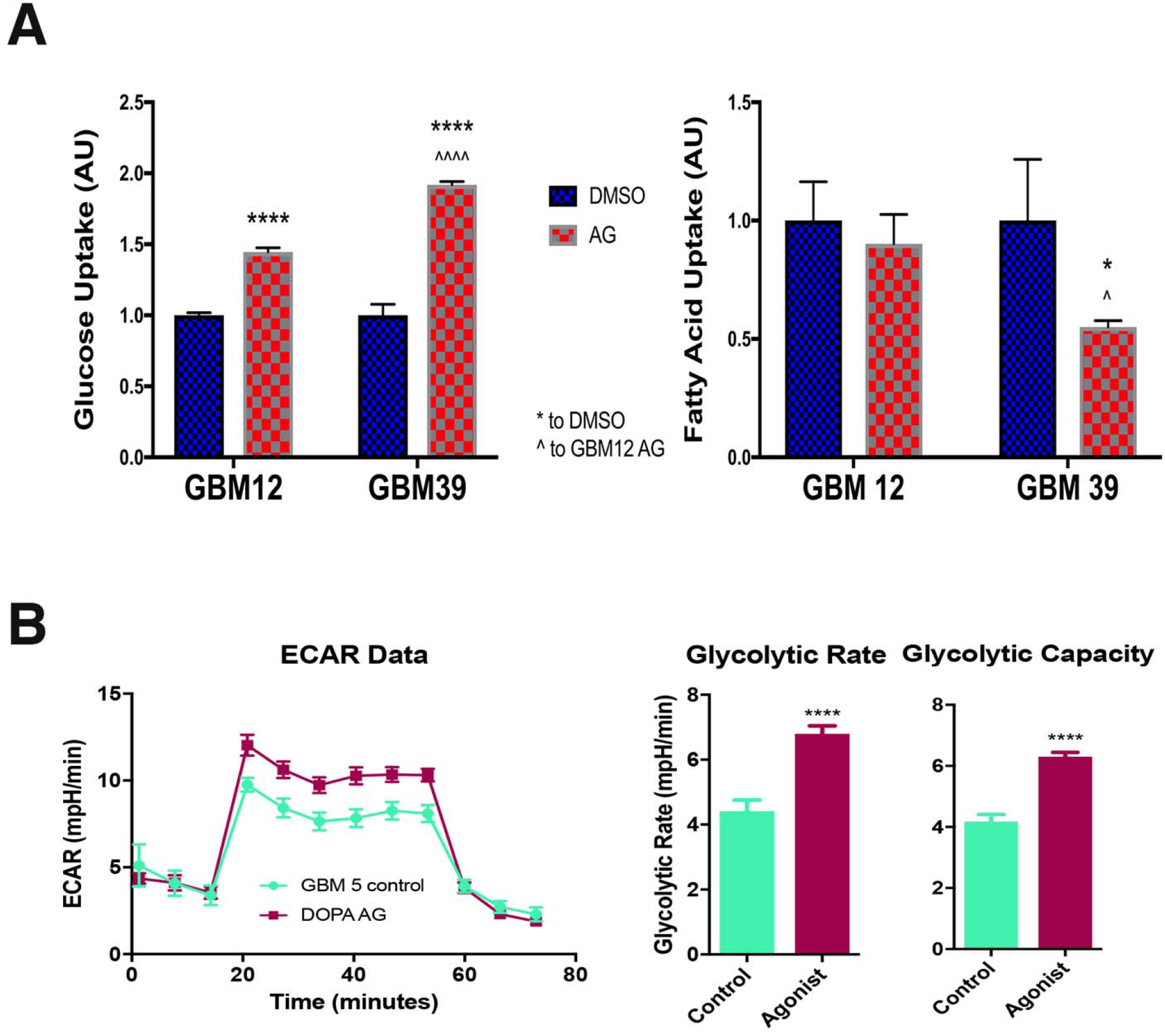
(A) Alkylating chemotherapy alters the uptake of glucose and fatty acid. PDX GBM cells were treated with either DMSO or TMZ. Four days later, cells were exposed to fluorescently-labeled glucose and palmitate. Uptake was determined by FACS analysis. (B) Seahorse analysis was used to determine the glycolytic rate of PDX lines after 4 days of 30nm agonist, or equimolar DMSO treatment. Comparisons were made using student t-Tests. *p<.05, **p<.01, ***p<.001.

## Acknowledgement

This work was supported by the National Institute of Neurological Disorders and Stroke grant 1R01NS096376, the American Cancer Society grant RSG-16-034-01-DDC (to A.U.A.) and National Cancer Institute grant R35CA197725 (to M.S.L) and P50CA221747 SPORE for Translational Approaches to Brain Cancer. We would like to thank Meijing Wu for the statistical analysis for the preparation of this manuscript.

